# Cleft lip/palate and educational attainment: cause, consequence, or correlation? A Mendelian randomization study

**DOI:** 10.1101/434126

**Authors:** Christina Dardani, Laurence J Howe, Evie Stergiakouli, Yvonne Wren, Kerry Humphries, Amy Davies, Karen Ho, Elisabeth Mangold, Kerstin U Ludwig, Caroline L Relton, George Davey Smith, Sarah J Lewis, Jonathan Sandy, Neil M Davies, Gemma C Sharp

**Affiliations:** Centre for Academic Mental Health, Population Health Sciences, Bristol Medical School, University of Bristol; Institute of Cardiovascular Science, University College London; MRC Integrative Epidemiology Unit, Population Health Sciences, Bristol Medical School, University of Bristol; The Cleft Collective, University of Bristol; MRC Integrative Epidemiology Unit, Bristol Dental School, University of Bristol; Bristol Speech and Language Therapy Research Unit, North Bristol NHS Trust; Bristol Bioresource Laboratories, Population Health Sciences, Bristol Medical School, University of Bristol; Institute of Human Genetics, University of Bonn; Department of Genomics, Life and Brain Center, University of Bonn; Dean of the Faculty of Health Sciences, University of Bristol

## Abstract

**Importance:** Previous studies have found that children born with a non-syndromic form of cleft lip and/or palate have lower-than-average educational attainment. These differences could be due to a genetic predisposition to low intelligence and academic performance, factors arising due to the cleft phenotype (such as school absence, social stigmatization and impaired speech and language development), or confounding by the prenatal environment. A clearer understanding of this mechanism will inform development of interventions to improve educational attainment in individuals born with a cleft, which could have wide-ranging knock-on effects on their quality of life.

**Objective:** To assess evidence for the hypothesis that common variant genetic liability to non-syndromic cleft lip with or without cleft palate (nsCL/P) influences educational attainment.

**Design:** Using summary data from genome-wide association studies (GWAS), we performed Linkage Disequilibrium (LD)-score regression and two-sample Mendelian randomization to evaluate the relationship between genetic liability to nsCL/P (GWAS n=3,987) and educational attainment (GWAS n=766,345), and intelligence (GWAS n=257,828).

**Results:** There was little evidence for shared genetic aetiology between nsCL/P and educational attainment (rg −0.03, 95% CI −0.14 to 0.08, P 0.58; βMR 0.002, 95% CI −0.001 to 0.005, P 0.417) or intelligence (rg −0.01, 95% CI −0.12 to 0.10, P 0.85; βMR 0.002, 95% CI −0.010 to 0.014, P 0.669).

**Conclusions and relevance:** Common genetic variants are unlikely to predispose individuals born with nsCL/P to low educational attainment or intelligence. This information will help tailor clinical-, school-, social- and family-level interventions to improve educational attainment in this group.

**Key Points:** *Question:* Do children born with a non-syndromic cleft lip with or without palate (nsCL/P) have lower-than average academic achievement because of an underlying genetic predisposition to educational attainment and/or intelligence?

*Findings:* There was little evidence for shared common variant genetic correlation between nsCL/P, educational attainment and intelligence.

*Meaning:* Common genetic variants are unlikely to predispose individuals born with nsCL/P to low educational attainment or intelligence. This information will help tailor clinical-, school-, social- and family-level interventions to improve educational attainment in this group.

## Introduction

Worldwide, orofacial clefts affect around one in 600-700 live births^1^. Although these structural anomalies can be surgically repaired (in regions where access to care is available), the condition remains associated with multiple adverse outcomes that can persist to adulthood, including impaired speech, appearance concerns and sub-optimal psychological wellbeing^2,3^.

Children born with orofacial clefts are at higher risk of low educational attainment, even when there are no other major birth defects or known syndromes. Small studies dating back to the 1950s have reported lower mean IQ scores, higher rates of learning difficulties and lower educational attainment in cases compared to controls or general population averages^4–9^. These findings have been corroborated by more recent, population-based studies. In a data linkage study in Atlanta, children with a cleft were three times more likely to use special education services than children with no major birth defects^10^. A Swedish population-based registry study showed that affected children (n=1,992) were less likely to receive high grades compared to over 1.2 million controls^11^. Similarly, studies based on registry data in Iowa showed that children with a non-syndromic cleft were approximately half a grade level behind their classmates^12^, with persistent low achievement trajectories^13^. Interestingly however, achievement scores were similar between affected children and their unaffected siblings^14^. In the most recent population-based study, 2,802 five-year-old children born with a non-syndromic cleft in England had lower average academic achievement across all learning domains compared to national averages^15^.

Low educational attainment can have a long-lasting adverse impact on vocational, social, mental and physical health outcomes^16^. Interventions and policies to improve educational attainment in individuals born with a cleft could have wide-ranging knock-on effects on their quality of life. However, it is currently unclear what the targets of such interventions should be, and indeed, whether these targets are even modifiable by intervention.

Therefore, we need to understand why individuals born with isolated, non-syndromic orofacial clefts have a higher risk of lower educational attainment. Three potential explanations for these associations are:

A. An underlying genetic liability to develop a cleft also influences intelligence and academic ability^17^, potentially via subtle undiagnosed congenital differences in brain structure or function^18,19^. Such effects could be caused by common or rare genetic variants. In previous work^20^, we have shown that common variant genetic liability to non-syndromic cleft influences facial morphology in individuals without a cleft, so a similar relationship might exist for educational attainment and intelligence.
B. Factors related to having a cleft influence educational attainment. Such factors include time spent under anaesthesia^21^, a high number of school absences due to healthcare appointments, social stigmatization (e.g. due to teasing by peers^22^, or perceptions and expectations of teachers^23^), lower self-esteem, or impaired speech^24^, or delayed language development^25^.
C. Environmental confounding by factors such as parental health behaviours or family socioeconomic status^14^ (Figure 1).

**Figure 1.**
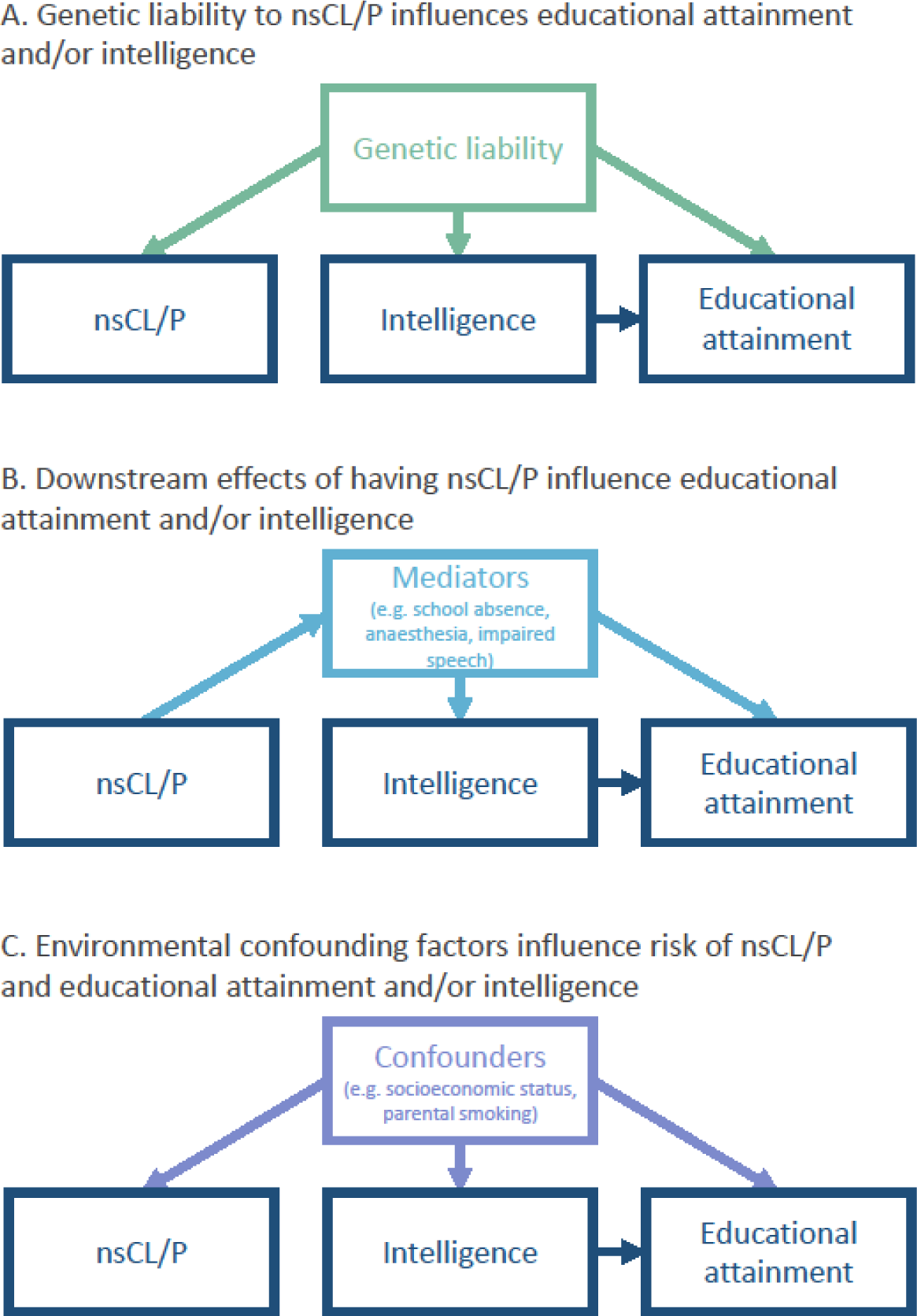
Potential explanations for observed associations between non-syndromic cleft lip with/without palate (nsCL/P) and lower educational attainment. In this study, we use genetic variants to test whether individuals born with nsCL/P are genetically predisposed to low educational attainment (Explanation A).

In this study, we used bidirectional Mendelian randomization (MR)^26,27^ and Linkage disequilibrium (LD) score regression^28^ to assess evidence for the hypothesis that genetic liability to non-syndromic cleft lip with or without cleft palate (nsCL/P), as captured by common genetic variation, influences educational attainment (Explanation A). A clearer understanding of this mechanism will help tailor interventions to improve educational attainment in individuals born with nsCL/P.

## Methods

We used LD score regression and MR to assess whether the association of nsCL/P and low educational attainment was due to genetic predisposition to low educational attainment or low intelligence. This analysis used summary statistics from published genome-wide association studies (GWAS).

### Samples (GWAS summary statistics)

#### Non-syndromic cleft lip/palate

For nsCL/P, we used a GWAS based on individual level genotype data from the International Cleft Consortium (ICC; dbGaP Study Accession phs000094.v1.p1), meta-analysed with GWAS summary statistics of the Bonn-II study^29^. The sample consisted of 1,215 nsCL/P cases and 2,772 parental and unrelated controls, all of European descent (total n=3,987). Further information on the generation of these GWAS statistics can be found in Howe et al., 2018^20^. This paper also shows that the summary statistics are comparable to those generated by a previous meta-analysis published by Ludwig et al.^30^, which used a similar approach in a sample of 666 European and European American trios and 795 Asian trios, combined with 399 cases and 1318 controls of European ancestry. Summary statistics from Ludwig et al. were not publicly available.

#### Educational attainment

For educational attainment, we used publicly available GWAS summary statistics published by Lee et al.^31^ (downloaded from https://www.thessgac.org/data), with a total sample size of 766,345 individuals. This was the total sample size available, excluding data from 23andMe due to restrictions on data sharing. Educational attainment was defined by mapping qualifications onto the International Standard of Classification of Education (ISCED) and was converted into years of education (in adults). This definition of educational attainment is strongly associated with other measures of educational attainment, including achieved grades and test scores^32^.

#### Intelligence

For intelligence, we used publicly available GWAS summary statistics published by Lee et al.^31^ (downloaded from https://www.thessgac.org/data), with a total sample size of 257,828 individuals. These summary statistics were generated by a meta-analysis of independent GWAS from UK Biobank and the COGENT consortium^33^. UK Biobank measured intelligence using a standardized score from a verbal-numerical reasoning test, designed as a measure of fluid intelligence. COGENT used a measure of intelligence based on performance on at least three neuropsychological tests or at least two IQ-test subscales. More information on phenotype definitions and generation of these GWAS summary statistics is available in Lee et al^31^.

### LD score regression

We used LD score regression to estimate the genetic correlation between liability to nsCL/P and both educational attainment and intelligence. LD score regression uses patterns of LD among genetic variants to estimate the extent of shared genetic aetiology among polygenic traits, accounting for cryptic relatedness and stratification^28^. We estimated genetic correlations using the suggested protocol for the LD score regression software for Python^28^, with pre-computed LD scores from the 1000 Genomes project^34^, available from the Broad Institute (https://data.broadinstitute.org/alkesgroup/LDSCORE/). In the regression analyses, we used an unconstrained intercept to account for (unknown, but unlikely) sample overlap.

### Bidirectional two-sample Mendelian randomization

#### The effect of liability to nsCL/P on educational attainment or intelligence

We applied two-sample summary statistic Mendelian randomization (MR) to assess whether liability to nsCL/P influences educational attainment and intelligence^26,27^. This approach enables estimation of causal effects from GWAS summary statistics. MR uses genetic variants (single nucleotide polymorphisms; SNPs) as proxies for the exposure that are not subject to confounding and reverse causation. The three main assumptions of MR are that i) SNPs are reliably associated with the exposure; ii) there are no confounders of the SNP-outcome association; and iii) the SNPs do not directly influence the outcome via a pathway independent of the exposure. The effect of the exposure on the outcome is calculated as the ratio of the SNP effect on the outcome by the effect of the SNP on the exposure. We conducted our two-sample MR analyses using the Two-Sample MR package for R^35^.

We selected 12 genome-wide significant SNPs as genetic instruments for nsCL/P from the nsCL/P GWAS meta-analysis published by Ludwig et al.^30^. This GWAS was conducted on a mixture of Europeans and Asians. We conducted a sensitivity analysis restricting to the six genome-wide significant SNPs identified in Europeans only. Summary statistics were not available for the study by Ludwig et al, so we extracted effect estimates and standard errors for the SNP-nsCL/P associations from the nsCL/P GWAS summary statistics described above and in Howe et al.^20^. SNP-outcome effect estimates and standard errors were extracted from the educational attainment and intelligence GWAS summary statistics described above.

Our primary analysis uses the inverse variance weighted (IVW) method. This method calculates the causal effect of genetic liability to nsCL/P (the exposure) on education/intelligence (the outcome) as the ratio of the SNP-outcome effect to the SNP-nsCL/P effect. We then conducted a series of sensitivity analyses to test the validity of the findings derived by the IVW approach. Specifically, we tested the consistency of our results to those obtained by: MR Egger^36^, weighted median^37^, and the weighted mode estimators^38^. MR Egger estimates the causal effect of the exposure on the outcome allowing for possible pleiotropic effects of the SNPs. The weighted median approach provides a causal effect estimate assuming that at least 50% of the SNPs in the analysis are valid instruments (i.e. the SNPs’ effect on the outcome is unconfounded and entirely mediated via the exposure). The weighted mode approach provides a causal estimate of the exposure on the outcome assuming the most common effect estimates come from SNPs that are valid instruments.

MR estimates and confidence intervals are expressed as a one unit increase in log odds of genetic liability to nsCL/P on standard deviations of years of education/IQ. To aid interpretation, we converted MR estimates into a scale describing the effect of a doubling in the genetic liability to nsCL/P on years of education or IQ points. To do this, we multiplied the original results by the standard deviation for the respective outcome (years of education SD = 3.8099, IQ SD = 15) as published by Lee et al.^31^. We then multiplied these figures by ln(2) to calculate the effect of a doubling of liability to nsCL/P.

#### The effect of educational attainment or intelligence on liability to nsCL/P

We also applied two sample MR in the reverse direction, i.e. to assess the causal effects of educational attainment and intelligence on offspring liability to nsCL/P. Since clefts form in the first ten weeks of embryonic development, any effect of education or intelligence will reflect parental effects – either due to passive transmission of parental genetics, or phenotypic expression of parental genetics that influences liability to nsCL/P in the offspring^39^. That is, nsCL/P cannot arise due to the child’s own education or intelligence, but parental genetic predisposition to low educational attainment or intelligence may influence the early prenatal environment to increase risk of nsCL/P^40^. Any parental effect can be inferred as being due to shared (50% from each parent) parent-offspring genetics^39^.

Of the 543,104 SNPs in the GWAS of nsCL/P, 522,190 were overlapping with the GWAS of educational attainment and 522,194 with the GWAS of intelligence. Of these overlapping SNPs, 2,346 and 1,018 had an effect allele frequency >=0.01 and a P-value <5*10^−8^ for the association with educational attainment or intelligence, respectively. After LD clumping, 368 approximately independent SNPs (r^2^=0.01, with a 10,000 kb window) were selected as instruments for educational attainment, and 132 for intelligence (Supplementary Table 3 and 4). We conducted IVW MR combining the SNP-educational attainment/intelligence and SNP-nsCL/P coefficients to give causal effect estimates of (parental) educational attainment and (parental) intelligence on (offspring) liability to nsCL/P, followed by sensitivity analyses.

MR estimates and confidence intervals are expressed as odds ratios for the effect of a one standard deviation unit increase in education/IQ on the odds of developing nsCL/P. To aid interpretation, we converted MR estimates into odds ratios for the effect of an extra year of education/an extra IQ point on the odds of developing nsCL/P. To do this, we converted to log odds and divided by the standard deviation for the respective traits (years of education SD = 3.8099, IQ SD = 15) as published by Lee et al.^31^. We then exponentiated these figures to convert to odds ratios.

### Data and code availability

All the data and code used to conduct the analyses described in this paper are provided in the Supplementary Material.

## Results

### Genetic correlation

Using LD score regression, we found little evidence of a substantial genetic correlation between liability to nsCL/P and educational attainment (rg −0.03, 95% CI −0.14 to 0.08, P 0.58) or intelligence (r_g_ −0.01, 95% CI −0.12 to 0.10, P 0.85).

### Mendelian randomization

Using bidirectional two sample MR, we found little evidence to suggest that genetic liability to nsCL/P influences educational attainment (IVW estimate 0.002; 95% CI −0.003 to 0.007; P 0.417). Although the MR estimate implies that a doubling in the genetic liability to nsCL/P increases years of education by 0.005 years, or around 1.9 days, the confidence interval crosses the null (−0.008 to 0.018 years of education per doubling in the genetic liability to nsCL/P). We also found little evidence for an effect of genetic liability to nsCL/P on intelligence (IVW estimate 0.002; 95% CI - 0.009 to 0.014; P 0.668). The MR estimate implies that a doubling in the genetic liability to nsCL/P increases intelligence by 0.02 IQ points, but again, the confidence interval crosses the null (−0.094 to 0.146 IQ points per doubling in the genetic liability to nsCL/P). These results were robust to sensitivity analyses using MR Egger, the weighted median and the weighted mode approach (Figure 2; Supplementary Tables 5-6). There was little evidence of horizontal pleiotropy bias in the causal estimate, as indicated by the MR Egger intercept (for educational attainment: 0.004, P 0.11; for intelligence: 0.008, P 0.14). Repeating our analysis restricting to the six genome-wide significant SNPs in the European only GWAS also did not change our findings (Supplementary Tables 7-8).

**Figure 2.**
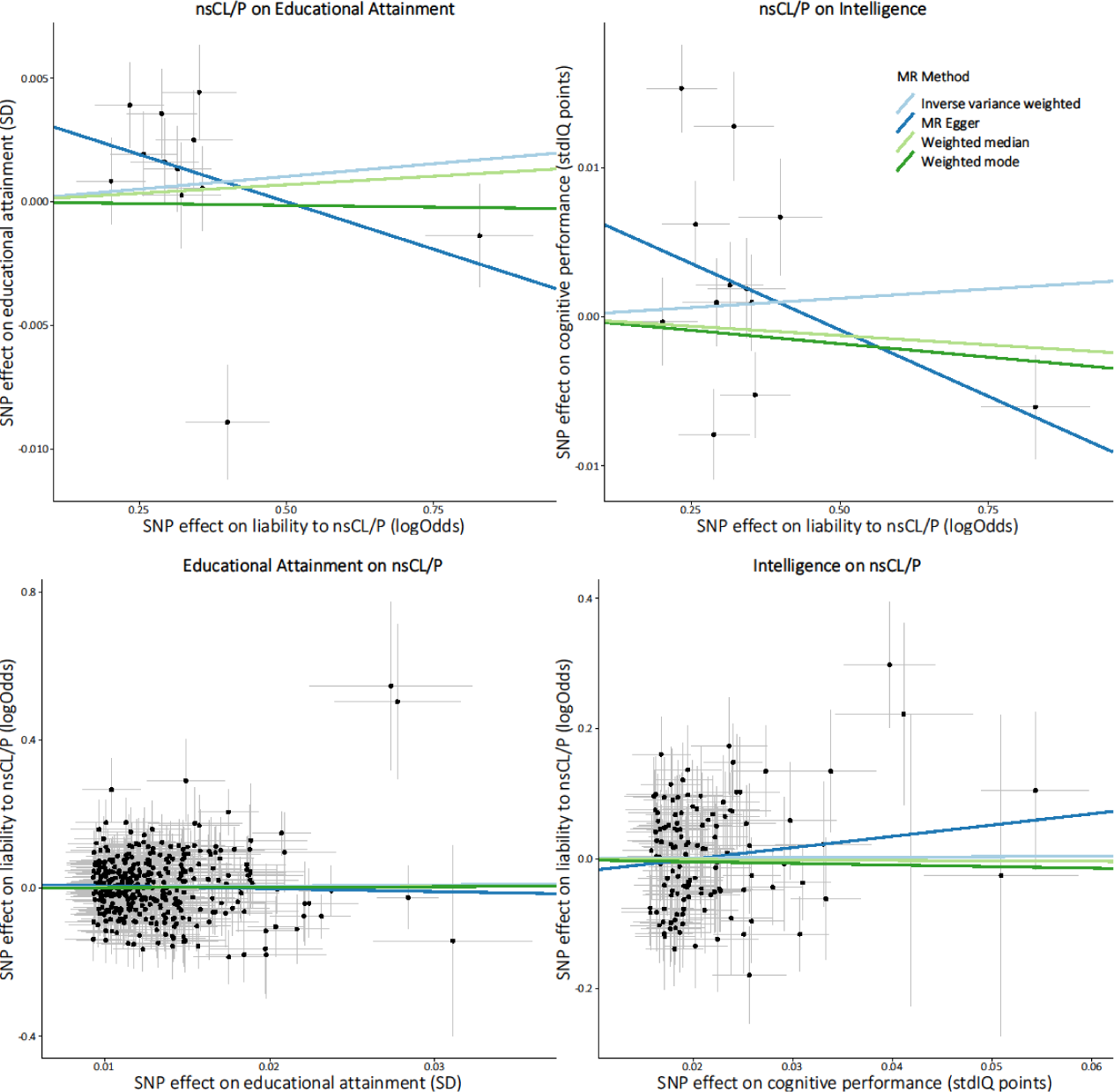
Bidirectional two sample Mendelian randomization results for associations between genetic liability to nsCL/P, educational attainment and intelligence, using four sensitivity analyses (inverse variance weighted, MR Egger, weighted median, and weighted mode).

We found little evidence of a causal effect of (parental) educational attainment on liability to nsCL/P (IVW odds ratio 1.34, 95% CI 0.75 to 2.38, P 0.32). Similarly, there was little evidence of a causal effect of (parental) intelligence on offspring liability to nsCL/P (IVW odds ratio 1.06, 95% CI 0.57 to 1.99, P 0.85) (Figure 2; Supplementary Tables 9-10).

## Discussion

### Summary of main findings

We found little evidence that educational attainment and intelligence were genetically correlated with, or affected by, genetic liability to nsCL/P. The large sample sizes in the GWASs of educational attainment and intelligence mean that this study was very well powered to detect an effect of nsCL/P, if it exists. This implies that individuals born with nsCL/P are unlikely to be genetically predisposed to spend less time in education or have lower intelligence (Explanation A in Figure 1). It seems more likely that the observed associations between nsCL/P and low educational attainment are explained by downstream, mediating factors related to being born with a cleft (such as time spent under anaesthesia, experience of bullying, impaired speech and delayed language development; Explanation B) or environmental confounding factors (such as socioeconomic position or parental health behaviours; Explanation C). This finding will help tailor interventions and policies that target factors influencing the observed associations to effectively improve educational attainment in this population.

### Comparison to previous evidence

In a previous study^20^, we found evidence that genetic liability to nsCL/P can influence facial morphology (specifically, philtrum width) in the general population, but the current study suggests it is unlikely that there is a similar relationship for educational attainment or intelligence.

There is evidence from the literature that nsCL/P is associated with downstream factors that might mediate any association between nsCL/P and educational attainment (Explanation B). Children born with a cleft, particularly involving the palate, are at higher risk of poor speech outcomes at three-years-old (i.e. before entering school) and persistent speech disorder^24^, both of which are strongly associated with lower educational attainment^41^. Teasing and bullying by peers is common in children born with clefts^22^, which can affect psychological wellbeing, enjoyment of school and attainment^42^. There is also some evidence that teachers perceive the behaviour and abilities of children born with a cleft differently from their classmates^23,43^. Affected children are required to take time off school to undergo surgery to repair the cleft (a study in the United States showed that roughly 24% of surgeries to repair CL and 37% of surgeries to repair CP are secondary surgeries, and roughly 70% of those occur during school ages^44^), and to attend follow-up health assessments, which could affect their learning. There is some observational evidence that repeated surgery (and therefore repeated exposure to general anaesthesia) is associated with lower IQ in children born with a cleft^21,45^.

There is also evidence suggesting that observed associations between nsCL/P and educational attainment might be explained by confounding (Explanation C). A registry-based study found similar levels of academic achievement in children with nsCL/P and their unaffected siblings^14^, which could indicate that any attainment deficit in children with nsCL/P is related to features of the family environment that are shared by unaffected family members. An alternative explanation for this finding is that the unaffected sibling is treated differently from the affected sibling in a way that reduces their educational attainment, for example, through divergence of parental attention and resources to the affected sibling.

Parental health behaviours, such as maternal smoking or alcohol consumption during pregnancy, have been linked to higher rates of nsCL/P^46^ and lower IQ and educational attainment in the general population^47,48^. In addition, many of the suggested risk factors for both nsCL/P and low educational attainment might be explained by confounding by lower family socioeconomic position, which has also been associated with nsCL/P^49^. In this study, we found little evidence for a causal effect of parental educational attainment on offspring nsCL/P. This does not support the hypothesis that familial socioeconomic position is a causal risk factor for nsCL/P. However, it should be noted that this interpretation is based on the assumptions that i) years of schooling is a good indication of socioeconomic position, ii) genetic variants in offspring are suitable instruments for parental educational attainment, and iii) the analysis was adequately powered to detect a clinically meaningful increase in risk.

### Strengths and limitations

The strengths of this study include: the novel application of a causal inference method (namely Mendelian randomization) to the effects of nsCL/P on education and intelligence; the use of non-overlapping samples drawn from the same population (European descent); the large sample sizes and statistical power to detect small effects in the LD score regression and MR analyses; the use of sensitivity analyses to test the robustness of our findings; and the publication of all the data and code used to conduct our analysis, which we hope will facilitate reproducibility and foster a culture of open science in cleft research.

There are also several factors that limit the interpretation of our findings: First, MR has several limitations (discussed in detail elsewhere^27,50,51^), such as horizontal pleiotropy which would violate one of the MR assumptions (when a SNP influences the outcome through a pathway other than via the exposure). We investigated this possibility using multiple independent genetic instruments as a sensitivity analysis (MR-Egger, the weighted median and the weighted mode approach). We found little evidence of pleiotropy. Furthermore, horizontal pleiotropy typically induces false positive findings, but is less likely to cause false negative results.

Second, we found little evidence of an effect of education on liability to nsCL/P. This could be because our estimates were not precise enough to detect the true causal effect. With only 3,987 samples (1,215 nsCL/P cases), this analysis had low power to detect modest effects.

Third, due to the design of the initial nsCL/P GWAS, which combined cleft lip only (CLO) with cleft lip with palate (CLP), we were unable to study subtype-specific effects, including any effect of cleft palate only (CPO), which was not studied in the GWAS we used. Findings from previous observational studies suggest that the orofacial cleft subtype is a strong predictor of academic outcomes^13^. Specifically, children with CPO are at higher risk of underperforming in several areas of academic learning, compared to both their unaffected peers, and also children born with CLO or CLP^52^. On the contrary, children born with CLO have been found to have academic achievement higher than children born with CLP or CPO^15^ and sometimes^53^ (though not always^13,15^) in-line with children born without a cleft.

Finally, because GWAS typically focus on common genetic variants, we were not able to investigate the potential contribution of rare genetic variants in explaining any shared genetic aetiology between nsCL/P, educational attainment and intelligence. High SNP heritability and low familial recurrence rates suggest that a substantial proportion of genetic liability to nsCL/P is likely to be captured by common genetic variation, but whole exome sequencing studies suggest that rare variants also contribute to the genetic aetiology of nsCL/P^54,55^. Furthermore, rare variants may cause syndromes involving CL/P, which could be misclassified as non-syndromic if the syndromes are difficult to identify clinically.

### Future work

This study highlights the need for further research to understand the multiple potential causes of lower educational attainment in individuals born with any type of orofacial cleft. Such research will require largescale, longitudinal data on affected children and their families, combining genetic data with detailed information on demographic, clinical, psychosocial, environmental, and developmental factors. The Cleft Collective Cohort Study^56,57^ was established in 2013 to address this need and help identify predictive and causal risk factors for cleft and cleft-related outcomes, including educational attainment. Its aim is to enable development of better strategies to facilitate early intervention to improve sub-optimal outcomes in individuals born with a cleft.

### Conclusion

This study shows that common genetic variants are unlikely to predispose individuals born with nsCL/P to low intelligence or educational attainment, highlighting the need for clinical-school-, social- and family-level interventions and policies to improve educational attainment in this population. The precise targets of these interventions will depend on further research to understand why individuals born with a cleft tend to do less well at school.

## Supporting information

## Funding acknowledgements

The Medical Research Council (MRC) and the University of Bristol support the MRC Integrative Epidemiology Unit [MC_UU_12013/1, MC_UU_12013/9, MC_UU_00011/1, MC_UU_00011/5]. The Scar Free Foundation supports the Cleft Collective (REC approval 13/SW/0064). The Economics and Social Research Council (ESRC) support NMD via a Future Research Leaders grant [ES/N000757/1]. CD is funded by the Wellcome Trust [108902/B/15/Z]. No funding body has influenced data collection, analysis or its interpretation. This publication is the work of the authors, who serve as the guarantors for the contents of this paper. This work was carried out using the computational facilities of the Advanced Computing Research Centre - http://www.bris.ac.uk/acrc/ and the Research Data Storage Facility of the University of Bristol - http://www.bris.ac.uk/acrc/storage/.

