## Supplementary material for "Cleft lip/palate and educational attainment: cause, consequence, or correlation? A Mendelian randomization study"

**Contents**

| Page |  |
| --- | --- |
| 2 | Supplementary Table 1: GWAS summary statistics used to conduct an MR of the effect of liability to nsCL/P on educational attainment. |
| 3 | Supplementary Table 2: GWAS summary statistics used to conduct an MR of the effect of liability to nsCL/P on intelligence. |
| 4 | Supplementary Table 3: GWAS summary statistics used to conduct an MR of the effect of educational attainment on liability to nsCL/P. |
| 12 | Supplementary Table 4: GWAS summary statistics used to conduct an MR of the effect of intelligence on liability to nsCL/P. |
| 15 | Supplementary Table 5: Causal effect estimates of genetic liability for nsCL/P on educational attainment derived by the primary MR analysis (IVW) as well as sensitivity analyses. |
| 16 | Supplementary Table 6: Causal effect estimates of genetic liability for nsCL/P on intelligence derived by the primary MR analysis (IVW) as well as sensitivity analyses. |
| 17 | Supplementary Table 7: Causal effect estimates of genetic liability for nsCL/P on educational attainment derived by the primary MR analysis (IVW) as well as sensitivity analyses. [RESTRICTING TO SNPS FROM EUROPEAN nsCL/P GWAS] |
| 18 | Supplementary Table 8: Causal effect estimates of genetic liability for nsCL/P on intelligence derived by the primary MR analysis (IVW) as well as sensitivity analyses. [RESTRICTING TO SNPS FROM EUROPEAN nsCL/P GWAS] |
| 19 | Supplementary Table 9: Causal effect estimates of educational attainment on nsCL/P derived by the primary MR analysis (IVW) as well as sensitivity analyses. |
| 20 | Supplementary Table 10: Causal effect estimates of intelligence on nsCL/P derived by the primary MR analysis (IVW) as well as sensitivity analyses. |
| 21 | Code used to run analyses |

Supplementary Table 1: GWAS summary statistics used to conduct an MR of the effect of liability to nsCL/P on educational attainment.

| SNP | Effect  Allele | Other Allele | nsCL/P Beta | nsCL/P SE | Education Beta | Education SE |
| --- | --- | --- | --- | --- | --- | --- |
| rs12543318 | A | C | -0.288 | 0.0594 | -0.007 | 0.003 |
| rs13041247 | T | C | 0.2018 | 0.0591 | 0.002 | 0.003 |
| rs1873147 | A | G | -0.3518 | 0.0621 | -0.001 | 0.003 |
| rs227731 | T | G | -0.3148 | 0.0564 | 0.000 | 0.002 |
| rs560426 | T | C | -0.2574 | 0.0564 | 0.000 | 0.002 |
| rs7078160 | A | G | 0.3996 | 0.0704 | -0.009 | 0.003 |
| rs742071 | T | G | 0.2927 | 0.0573 | 0.003 | 0.003 |
| rs7590268 | T | G | -0.3428 | 0.065 | -0.001 | 0.003 |
| rs7632427 | T | C | 0.2336 | 0.0583 | 0.003 | 0.003 |
| rs8001641 | A | G | 0.3573 | 0.0581 | 0.000 | 0.002 |
| rs861020 | A | G | 0.3218 | 0.066 | 0.004 | 0.003 |
| rs987525 | A | C | 0.8286 | 0.0909 | -0.001 | 0.003 |

Supplementary Table 2: GWAS summary statistics used to conduct an MR of the effect of liability to nsCL/P on intelligence.

| SNP | Effect Allele | Other Allele | nsCL/P Beta | nsCL/P SE | Intelligence Beta | Intelligence SE |
| --- | --- | --- | --- | --- | --- | --- |
| rs12543318 | A | C | -0.288 | 0.0594 | 0.00789 | 0.00301 |
| rs13041247 | T | C | 0.2018 | 0.0591 | -0.00034 | 0.00291 |
| rs1873147 | A | G | -0.3518 | 0.0621 | -0.00097 | 0.0032 |
| rs227731 | T | G | -0.3148 | 0.0564 | -0.00212 | 0.00288 |
| rs560426 | T | C | -0.2574 | 0.0564 | -0.0062 | 0.00286 |
| rs7078160 | A | G | 0.3996 | 0.0704 | 0.0067 | 0.00388 |
| rs742071 | T | G | 0.2927 | 0.0573 | 0.00096 | 0.00292 |
| rs7590268 | T | G | -0.3428 | 0.065 | -0.00189 | 0.00334 |
| rs7632427 | T | C | 0.2336 | 0.0583 | 0.01532 | 0.00291 |
| rs8001641 | A | G | 0.3573 | 0.0581 | -0.00522 | 0.00285 |
| rs861020 | A | G | 0.3218 | 0.066 | 0.01277 | 0.0036 |
| rs987525 | A | C | 0.8286 | 0.0909 | -0.00605 | 0.00346 |

Supplementary Table 3: GWAS summary statistics used to conduct an MR of the effect of educational attainment on liability to nsCL/P.

| SNP | Effect Allele | Other Allele | Education Beta | Education SE | nsCL/P Beta | nsCL/P SE |
| --- | --- | --- | --- | --- | --- | --- |
| rs10072007 | T | C | -0.01535 | 0.00253 | 0.0247 | 0.1275 |
| rs10079990 | A | C | 0.00978 | 0.00178 | -0.0126 | 0.0583 |
| rs10086223 | C | T | -0.01002 | 0.00172 | -0.0363 | 0.0587 |
| rs1008661 | A | G | 0.01375 | 0.00181 | -0.0011 | 0.0609 |
| rs1014174 | C | T | -0.00933 | 0.0017 | -0.0296 | 0.0563 |
| rs10180461 | C | T | 0.01627 | 0.00175 | 0.0248 | 0.0589 |
| rs10184774 | C | T | 0.01082 | 0.00183 | -0.0153 | 0.0611 |
| rs10201404 | T | G | 0.02239 | 0.00282 | -0.0406 | 0.0928 |
| rs10205801 | A | G | -0.01053 | 0.00171 | 0.0153 | 0.0571 |
| rs10236197 | C | T | -0.01161 | 0.00177 | -0.0488 | 0.0602 |
| rs1025206 | G | A | -0.01099 | 0.002 | -0.0319 | 0.0662 |
| rs10270472 | G | A | -0.01022 | 0.00171 | 0.0144 | 0.0567 |
| rs10416924 | A | G | -0.01008 | 0.00174 | 0.0036 | 0.0575 |
| rs10448005 | T | C | -0.01507 | 0.0026 | -0.0929 | 0.0851 |
| rs10483837 | A | G | -0.0145 | 0.00211 | -0.1151 | 0.0712 |
| rs10484022 | A | G | 0.01035 | 0.0017 | -0.0161 | 0.0567 |
| rs10490175 | A | G | -0.01465 | 0.00217 | -0.0628 | 0.0712 |
| rs10496347 | G | A | 0.00996 | 0.00174 | 0.0147 | 0.0815 |
| rs10510388 | C | A | 0.01148 | 0.00177 | 0.02 | 0.0594 |
| rs10511271 | T | C | -0.01392 | 0.00202 | -0.0874 | 0.0678 |
| rs10513524 | A | G | 0.01979 | 0.00363 | -0.1797 | 0.1181 |
| rs10513656 | G | A | 0.01479 | 0.0021 | -0.1312 | 0.0688 |
| rs10515086 | C | T | 0.01478 | 0.00231 | 0.1053 | 0.0787 |
| rs10516930 | T | C | -0.01089 | 0.00187 | 0.02 | 0.0623 |
| rs10517967 | G | A | 0.01125 | 0.0019 | 0.0701 | 0.0616 |
| rs10748815 | A | G | -0.01675 | 0.00172 | -0.1121 | 0.0571 |
| rs10760193 | T | C | 0.01438 | 0.00172 | -0.0057 | 0.0583 |
| rs10761247 | A | G | 0.00995 | 0.00173 | -0.0333 | 0.0572 |
| rs10765777 | A | C | -0.01477 | 0.00176 | -0.0181 | 0.058 |
| rs1077084 | G | A | 0.00978 | 0.0017 | -0.0326 | 0.0566 |
| rs10805383 | A | G | -0.01043 | 0.0017 | 0.0215 | 0.0567 |
| rs10810098 | T | C | -0.01482 | 0.0019 | 0.0117 | 0.0648 |
| rs10818288 | G | T | 0.01077 | 0.0017 | -0.056 | 0.0565 |
| rs10822736 | T | C | 0.01133 | 0.00172 | -0.0072 | 0.0583 |
| rs10845987 | T | C | -0.01316 | 0.00171 | 0.0343 | 0.0567 |
| rs10854884 | C | A | -0.01467 | 0.00172 | -0.0593 | 0.0577 |
| rs10872224 | T | G | -0.02314 | 0.00174 | 0.076 | 0.0586 |
| rs10887801 | G | T | -0.01087 | 0.00171 | 0.0562 | 0.0567 |
| rs10915206 | G | T | 0.01079 | 0.00187 | 0.0871 | 0.0644 |
| rs10922911 | T | C | -0.01658 | 0.00175 | 0.0646 | 0.0582 |
| rs11012 | C | T | 0.01846 | 0.00225 | -0.1791 | 0.0744 |
| rs11048916 | C | T | 0.01049 | 0.00178 | 0.0247 | 0.0596 |
| rs1106090 | A | G | 0.01173 | 0.00175 | -0.0172 | 0.0586 |
| rs11071848 | T | C | -0.01443 | 0.00224 | 0.046 | 0.0758 |
| rs1108343 | T | C | -0.01084 | 0.00179 | -0.0423 | 0.0598 |
| rs11115011 | A | G | 0.01493 | 0.00234 | 0.2906 | 0.1107 |
| rs11157930 | G | T | 0.01361 | 0.00174 | -0.0723 | 0.083 |
| rs11160755 | G | T | 0.01157 | 0.00193 | 0.0117 | 0.0882 |
| rs11210869 | A | G | -0.01787 | 0.00176 | -0.1051 | 0.0592 |
| rs11223560 | G | A | -0.00995 | 0.00173 | -0.1334 | 0.0588 |
| rs11243852 | T | C | 0.01149 | 0.00199 | 0.093 | 0.0671 |
| rs11564410 | A | G | 0.01126 | 0.00191 | 0.0406 | 0.0648 |
| rs11588857 | G | A | -0.02208 | 0.00208 | 0.0756 | 0.0689 |
| rs11623285 | G | T | 0.01413 | 0.00251 | -0.1572 | 0.0876 |
| rs11635966 | C | T | 0.01228 | 0.00177 | -0.0469 | 0.0585 |
| rs11644573 | C | T | -0.00969 | 0.00171 | -0.0494 | 0.057 |
| rs11665242 | A | G | 0.01297 | 0.00171 | 0.0166 | 0.0558 |
| rs1167827 | G | A | -0.00969 | 0.00173 | 0.0139 | 0.0574 |
| rs11689426 | T | C | 0.01425 | 0.00258 | -0.1034 | 0.0862 |
| rs11704277 | C | T | -0.01548 | 0.00284 | -0.1754 | 0.0965 |
| rs11706370 | G | A | -0.02838 | 0.00184 | 0.0255 | 0.0852 |
| rs11720121 | T | C | -0.01386 | 0.00173 | -0.0698 | 0.0584 |
| rs11724252 | A | G | -0.01352 | 0.0018 | 0.0206 | 0.0606 |
| rs11764590 | T | C | 0.01753 | 0.00215 | -0.1861 | 0.072 |
| rs11772832 | T | C | 0.01213 | 0.00174 | -0.1274 | 0.0586 |
| rs11782215 | A | G | -0.0158 | 0.00175 | 0.0574 | 0.0578 |
| rs11849844 | T | C | 0.02774 | 0.00381 | 0.5032 | 0.2092 |
| rs11871429 | A | G | 0.01425 | 0.00202 | -0.0426 | 0.0982 |
| rs11901438 | C | T | 0.01156 | 0.00185 | 0.0504 | 0.0623 |
| rs11954686 | A | G | 0.01241 | 0.00221 | 0.1433 | 0.0743 |
| rs12024263 | G | A | -0.01698 | 0.00203 | -0.0799 | 0.069 |
| rs12170452 | G | A | -0.01153 | 0.00171 | -0.026 | 0.0572 |
| rs12192936 | A | G | 0.01207 | 0.00201 | 0.0331 | 0.0674 |
| rs12259847 | C | T | 0.01194 | 0.00218 | 0.0257 | 0.1016 |
| rs12342546 | G | A | 0.01126 | 0.00203 | 0.068 | 0.0644 |
| rs12569425 | G | A | -0.01514 | 0.00249 | 0.0833 | 0.0828 |
| rs12576777 | C | T | -0.01109 | 0.00177 | -0.0098 | 0.0588 |
| rs12588988 | C | T | 0.01053 | 0.0017 | 0.0433 | 0.0567 |
| rs12598842 | G | A | 0.00969 | 0.00174 | -0.0323 | 0.0585 |
| rs12610468 | T | C | -0.01098 | 0.0017 | 0.0557 | 0.0574 |
| rs12646523 | C | T | 0.01185 | 0.00196 | 0.0049 | 0.066 |
| rs1266416 | T | C | 0.01084 | 0.00188 | 0.009 | 0.0617 |
| rs12670376 | G | A | -0.01062 | 0.00171 | 0.0654 | 0.0568 |
| rs1267501 | C | T | -0.01775 | 0.00219 | 0.0542 | 0.0734 |
| rs12741781 | T | G | 0.01475 | 0.00181 | -0.0798 | 0.0609 |
| rs12768534 | G | A | 0.01361 | 0.0017 | 0.0156 | 0.0576 |
| rs12789313 | T | C | 0.01001 | 0.0017 | -0.0676 | 0.0822 |
| rs12806862 | G | A | -0.00998 | 0.00172 | -0.0385 | 0.0576 |
| rs1291818 | C | T | -0.01085 | 0.0017 | 0.06 | 0.0562 |
| rs12934778 | C | T | 0.01365 | 0.00242 | -0.0237 | 0.0818 |
| rs12965322 | A | C | 0.00977 | 0.0017 | 0.0315 | 0.0561 |
| rs13010010 | C | T | -0.02071 | 0.00175 | -0.1483 | 0.0591 |
| rs13094951 | G | A | 0.02214 | 0.00324 | -0.0415 | 0.1115 |
| rs13107325 | C | T | 0.0188 | 0.00344 | 0.1045 | 0.121 |
| rs13160109 | G | A | -0.0121 | 0.00211 | -0.049 | 0.0699 |
| rs13210693 | A | G | 0.00932 | 0.0017 | -0.1376 | 0.0573 |
| rs1322537 | T | C | -0.01829 | 0.00225 | 0.0131 | 0.0755 |
| rs1328687 | A | C | -0.00978 | 0.0017 | -0.0297 | 0.0569 |
| rs1329044 | C | T | -0.01437 | 0.00224 | 0.0961 | 0.0732 |
| rs13400118 | A | C | 0.01385 | 0.0018 | -0.065 | 0.0827 |
| rs13422673 | C | T | 0.01201 | 0.0017 | 0.0101 | 0.0564 |
| rs13424809 | A | G | -0.03109 | 0.00482 | 0.1424 | 0.2576 |
| rs13429247 | A | G | -0.01418 | 0.00192 | 0.1349 | 0.0629 |
| rs1343700 | A | G | 0.01026 | 0.00177 | -0.0634 | 0.084 |
| rs1358596 | C | A | 0.01415 | 0.00235 | -0.0836 | 0.0758 |
| rs1360382 | A | G | -0.01864 | 0.00182 | -0.05 | 0.0609 |
| rs1362910 | G | A | 0.01204 | 0.00174 | 0.0664 | 0.0575 |
| rs1371638 | T | C | 0.01295 | 0.00183 | 0.0552 | 0.0601 |
| rs1376503 | G | A | 0.01177 | 0.00191 | -0.1519 | 0.0639 |
| rs1391438 | T | C | 0.0167 | 0.00183 | -0.0643 | 0.0603 |
| rs1400875 | T | C | 0.01093 | 0.0018 | -0.0667 | 0.0848 |
| rs140147 | A | G | -0.0112 | 0.00176 | 0.0934 | 0.0592 |
| rs1425542 | C | T | -0.01672 | 0.00209 | -0.0958 | 0.0694 |
| rs1427298 | C | T | -0.0102 | 0.00172 | -0.1012 | 0.0583 |
| rs1439984 | T | C | -0.00968 | 0.00172 | -0.1584 | 0.0798 |
| rs1449172 | T | G | 0.01253 | 0.00186 | 0.0667 | 0.063 |
| rs1449477 | A | C | 0.0095 | 0.0017 | 0.0407 | 0.0565 |
| rs1450782 | G | T | -0.00945 | 0.00173 | -0.0623 | 0.0588 |
| rs1452075 | T | C | 0.01226 | 0.00193 | -0.0317 | 0.0648 |
| rs1465900 | C | A | 0.01264 | 0.00208 | 0.0217 | 0.0672 |
| rs1477031 | G | A | -0.00976 | 0.00174 | -0.0693 | 0.0583 |
| rs1477114 | C | T | 0.01055 | 0.0017 | -0.019 | 0.0577 |
| rs1479679 | C | T | -0.01704 | 0.00263 | -0.1024 | 0.0886 |
| rs1529127 | A | G | 0.0097 | 0.0017 | 0.0154 | 0.0563 |
| rs1550816 | C | T | -0.01107 | 0.00172 | -0.0115 | 0.0565 |
| rs1555366 | T | C | 0.01199 | 0.002 | -0.0087 | 0.0665 |
| rs1567130 | A | C | 0.01063 | 0.00185 | 0.0574 | 0.0876 |
| rs1567540 | A | G | -0.01271 | 0.00219 | -0.0743 | 0.0724 |
| rs1569092 | A | G | 0.01807 | 0.00234 | -0.0373 | 0.0802 |
| rs1569723 | A | C | 0.01168 | 0.00197 | 0.1116 | 0.0666 |
| rs1599389 | A | G | 0.00979 | 0.0017 | 0.0136 | 0.0566 |
| rs1612548 | G | A | 0.01475 | 0.00204 | 0.0494 | 0.0698 |
| rs1620977 | G | A | -0.02046 | 0.00195 | 0.0026 | 0.0646 |
| rs1671476 | A | G | -0.01882 | 0.00323 | -0.0453 | 0.1115 |
| rs16843629 | A | G | -0.01237 | 0.00184 | -0.001 | 0.0606 |
| rs16854920 | T | C | -0.01007 | 0.00181 | -0.0028 | 0.0604 |
| rs16960499 | G | A | -0.01708 | 0.00264 | 0.0401 | 0.0867 |
| rs1701 | C | A | -0.01262 | 0.00191 | -0.076 | 0.0832 |
| rs1701704 | T | G | -0.0175 | 0.0018 | -0.2052 | 0.0601 |
| rs1704147 | T | C | -0.01249 | 0.00183 | 0.0302 | 0.0618 |
| rs17119973 | G | A | 0.01402 | 0.00194 | 0.1124 | 0.0669 |
| rs17123764 | T | C | 0.0187 | 0.0033 | 0.035 | 0.1049 |
| rs1734394 | T | C | -0.01153 | 0.00183 | 0.0588 | 0.0616 |
| rs17428262 | C | T | -0.01975 | 0.00341 | 0.1637 | 0.1203 |
| rs17552299 | G | A | 0.02734 | 0.00493 | 0.5463 | 0.2272 |
| rs17565975 | A | G | -0.01142 | 0.00171 | -0.021 | 0.0571 |
| rs176218 | G | T | -0.01883 | 0.00215 | -0.0411 | 0.0961 |
| rs17770697 | G | A | -0.01376 | 0.002 | -0.0782 | 0.0895 |
| rs17785248 | G | A | -0.01216 | 0.00198 | -0.0434 | 0.0668 |
| rs1783978 | C | T | -0.00951 | 0.00171 | -0.1216 | 0.0557 |
| rs1829534 | A | G | -0.00967 | 0.00172 | -0.0646 | 0.0572 |
| rs1866823 | A | G | 0.01009 | 0.00171 | -0.1151 | 0.0809 |
| rs1880692 | A | G | 0.01008 | 0.0017 | -0.0196 | 0.0569 |
| rs1891611 | G | A | -0.00972 | 0.00173 | 0.0691 | 0.0582 |
| rs1896297 | C | T | 0.01486 | 0.0018 | 0.0188 | 0.0604 |
| rs191092 | C | T | -0.01634 | 0.00296 | 0.0915 | 0.1007 |
| rs1913332 | T | G | 0.00969 | 0.00174 | 0.0588 | 0.0579 |
| rs1920045 | T | C | 0.00961 | 0.00173 | -0.0995 | 0.0584 |
| rs1938570 | T | C | -0.01862 | 0.00232 | -0.0635 | 0.0758 |
| rs1962299 | A | G | 0.00961 | 0.00176 | 0.0413 | 0.0582 |
| rs1964927 | G | A | -0.01423 | 0.00177 | 0.0812 | 0.0586 |
| rs1977552 | T | G | -0.01156 | 0.00195 | -0.0434 | 0.0659 |
| rs1982564 | T | G | 0.00935 | 0.00171 | -0.0914 | 0.0572 |
| rs1998086 | G | A | -0.01369 | 0.00208 | 0.1245 | 0.0663 |
| rs2002058 | C | T | 0.01211 | 0.00216 | -0.0181 | 0.0713 |
| rs2003154 | A | G | 0.0169 | 0.00262 | 0.0213 | 0.0872 |
| rs2014830 | C | T | -0.01049 | 0.00186 | -0.0772 | 0.061 |
| rs2054122 | T | G | 0.01543 | 0.0017 | 0.0255 | 0.0577 |
| rs2058943 | T | C | -0.01027 | 0.00184 | -0.0058 | 0.0628 |
| rs2068428 | T | C | 0.01895 | 0.00198 | 0.0132 | 0.0656 |
| rs2164300 | C | T | -0.01282 | 0.00171 | -0.0401 | 0.0571 |
| rs2179152 | C | T | 0.01455 | 0.00176 | 0.0253 | 0.058 |
| rs2182505 | T | C | 0.01086 | 0.00192 | 0.0403 | 0.0635 |
| rs2202769 | G | A | -0.01472 | 0.00242 | -0.0771 | 0.0796 |
| rs2216144 | C | T | -0.01043 | 0.0017 | -0.0216 | 0.0568 |
| rs2252641 | T | C | -0.00966 | 0.00171 | 0.0561 | 0.0577 |
| rs2256965 | G | A | -0.01128 | 0.00176 | 0.0773 | 0.0579 |
| rs2269933 | G | A | 0.0102 | 0.00183 | -0.0175 | 0.0632 |
| rs2281767 | T | C | -0.0101 | 0.0017 | 0.089 | 0.0573 |
| rs2283076 | A | G | 0.01143 | 0.00204 | 0.0102 | 0.0983 |
| rs2291725 | T | C | 0.01033 | 0.00172 | 0.077 | 0.0571 |
| rs2295709 | C | T | 0.01373 | 0.00208 | -0.0317 | 0.0715 |
| rs2298679 | T | C | -0.0118 | 0.00208 | 0.0894 | 0.0691 |
| rs2302761 | T | C | 0.01354 | 0.00209 | -0.0262 | 0.0719 |
| rs2307377 | A | G | 0.0209 | 0.0031 | 0.0971 | 0.1073 |
| rs2333256 | G | T | 0.01188 | 0.0017 | -0.091 | 0.0568 |
| rs235509 | G | T | -0.01349 | 0.00222 | -0.1238 | 0.0751 |
| rs242093 | A | G | -0.01031 | 0.00172 | -0.0461 | 0.0571 |
| rs2447540 | G | A | -0.01176 | 0.00184 | -0.035 | 0.0607 |
| rs2529044 | T | C | 0.01877 | 0.00218 | 0.0362 | 0.0737 |
| rs252991 | G | A | -0.00998 | 0.00177 | 0.0359 | 0.0593 |
| rs2544809 | C | T | -0.01294 | 0.00172 | -0.1065 | 0.0573 |
| rs2562779 | G | A | -0.0112 | 0.00202 | 0.0164 | 0.0677 |
| rs2570497 | C | T | 0.01233 | 0.00177 | -0.1651 | 0.0597 |
| rs2588917 | A | C | -0.00947 | 0.00171 | 0.0095 | 0.0578 |
| rs25934 | G | T | 0.01489 | 0.00246 | -0.1416 | 0.0854 |
| rs2610225 | A | G | 0.0104 | 0.0017 | -0.0171 | 0.056 |
| rs2664299 | C | T | 0.01042 | 0.00172 | 0.0452 | 0.0571 |
| rs268134 | G | A | -0.01235 | 0.00196 | 0.0267 | 0.0639 |
| rs2706762 | T | C | 0.01484 | 0.00246 | 0.082 | 0.0801 |
| rs2718794 | T | C | -0.01503 | 0.00258 | -0.1113 | 0.0881 |
| rs2721195 | T | C | -0.01142 | 0.00171 | -0.013 | 0.0573 |
| rs273438 | G | A | 0.00937 | 0.00171 | 0.0961 | 0.0571 |
| rs277828 | C | A | 0.01091 | 0.00196 | 0.0467 | 0.067 |
| rs2787101 | C | T | -0.00968 | 0.00174 | -0.0731 | 0.0585 |
| rs2833476 | T | C | -0.01886 | 0.0034 | -0.1291 | 0.1534 |
| rs2852714 | A | G | 0.01337 | 0.00192 | 0.09 | 0.0636 |
| rs2906457 | A | C | -0.01238 | 0.00194 | -0.1244 | 0.0644 |
| rs2910606 | T | G | -0.01143 | 0.0017 | 0.0304 | 0.057 |
| rs2971970 | T | G | -0.01654 | 0.00207 | 0.0274 | 0.0703 |
| rs2974298 | C | T | -0.01077 | 0.0017 | -0 | 0.057 |
| rs298601 | C | T | 0.01066 | 0.0017 | 0.0278 | 0.0565 |
| rs2997063 | T | C | -0.01006 | 0.00172 | -0.0009 | 0.0575 |
| rs301788 | T | C | -0.01483 | 0.00225 | -0.0927 | 0.0727 |
| rs3103472 | T | C | -0.01128 | 0.00178 | -0.0303 | 0.0595 |
| rs3131576 | T | C | 0.01003 | 0.00171 | 0.0546 | 0.0565 |
| rs321241 | A | G | -0.01306 | 0.00173 | 0.0327 | 0.0578 |
| rs339054 | G | T | 0.0117 | 0.0017 | 0.0527 | 0.0576 |
| rs34320 | G | T | -0.01977 | 0.00177 | 0.1156 | 0.0582 |
| rs363096 | C | T | 0.01363 | 0.00172 | -0.0143 | 0.0583 |
| rs3738773 | G | A | -0.01276 | 0.00186 | 0.0431 | 0.0625 |
| rs3769941 | A | G | -0.01061 | 0.00193 | -0.0055 | 0.0649 |
| rs3785354 | T | C | -0.01737 | 0.00176 | -0.0114 | 0.0579 |
| rs3794621 | G | A | 0.01113 | 0.00172 | -0.083 | 0.0813 |
| rs3796627 | T | C | 0.01358 | 0.00178 | 0.0009 | 0.0595 |
| rs37976 | T | C | 0.01068 | 0.00171 | -0.0534 | 0.0577 |
| rs3935685 | C | T | 0.01265 | 0.00171 | 0.0374 | 0.0567 |
| rs4073894 | A | G | 0.01524 | 0.00211 | 0.0497 | 0.1053 |
| rs4074838 | G | A | 0.01354 | 0.00215 | -0.0074 | 0.0703 |
| rs4129585 | A | C | -0.0114 | 0.00171 | 0.0027 | 0.0577 |
| rs4130548 | T | C | 0.00976 | 0.00176 | 0.1024 | 0.0595 |
| rs4144624 | C | T | 0.01338 | 0.00239 | -0.0467 | 0.0826 |
| rs4235481 | C | T | -0.02036 | 0.0017 | 0.1041 | 0.0811 |
| rs4236176 | G | A | -0.01159 | 0.00195 | -0.1519 | 0.0654 |
| rs4240927 | T | C | 0.00988 | 0.0017 | -0.0051 | 0.0566 |
| rs4240966 | A | C | 0.0116 | 0.00178 | -0.0503 | 0.06 |
| rs4265116 | G | T | 0.01463 | 0.0025 | 0.0345 | 0.0821 |
| rs4353689 | C | A | -0.0182 | 0.00251 | -0.0011 | 0.0831 |
| rs4363310 | T | C | -0.01544 | 0.00183 | -0.1152 | 0.0616 |
| rs4377285 | A | G | 0.01315 | 0.00225 | -0.0944 | 0.0747 |
| rs4383040 | T | C | 0.01166 | 0.00178 | 0.0839 | 0.0855 |
| rs4441609 | T | C | -0.01042 | 0.00175 | -0.0744 | 0.0582 |
| rs4455149 | A | G | -0.01118 | 0.00179 | -0.0278 | 0.0593 |
| rs4580739 | G | A | 0.01289 | 0.00189 | -0.0801 | 0.0623 |
| rs4585573 | G | T | -0.01027 | 0.00178 | -0.0203 | 0.0601 |
| rs4600366 | A | G | -0.01178 | 0.0019 | -0.0636 | 0.0619 |
| rs4695140 | T | C | -0.01037 | 0.00175 | 0.0843 | 0.0602 |
| rs4697414 | T | C | -0.0158 | 0.00227 | -0.1007 | 0.1122 |
| rs4726070 | G | A | -0.01251 | 0.00174 | 0.084 | 0.0576 |
| rs4757956 | A | G | 0.01372 | 0.00184 | 0.053 | 0.0615 |
| rs4781236 | T | C | 0.01041 | 0.00171 | -0.0516 | 0.057 |
| rs4782262 | T | C | 0.01734 | 0.00283 | -0.0677 | 0.091 |
| rs4799088 | A | G | 0.01193 | 0.00189 | 0.1205 | 0.0648 |
| rs4832633 | G | A | 0.0112 | 0.00192 | 0.0763 | 0.0623 |
| rs4835349 | C | T | 0.01251 | 0.00186 | -0.0265 | 0.0619 |
| rs4836131 | C | T | -0.01261 | 0.00196 | -0.0096 | 0.0659 |
| rs4839154 | C | A | 0.01243 | 0.002 | 0.0339 | 0.0659 |
| rs4851758 | T | C | -0.0141 | 0.00257 | 0.1296 | 0.0841 |
| rs4858670 | C | T | 0.0101 | 0.00183 | -0.0075 | 0.0607 |
| rs4877152 | T | G | 0.01157 | 0.00171 | -0.0013 | 0.0571 |
| rs4904523 | G | A | 0.00936 | 0.0017 | -0.0145 | 0.0564 |
| rs4906679 | T | C | 0.00946 | 0.0017 | 0.0676 | 0.0568 |
| rs4925109 | A | G | -0.01075 | 0.00184 | 0.1206 | 0.084 |
| rs4958573 | A | G | -0.01115 | 0.00191 | 0.0875 | 0.062 |
| rs4972551 | G | T | -0.01149 | 0.00181 | 0.0428 | 0.0887 |
| rs4974424 | A | G | -0.0149 | 0.00234 | -0.0445 | 0.0787 |
| rs4976447 | A | G | 0.0119 | 0.00197 | -0.0587 | 0.0649 |
| rs4984683 | A | C | -0.0117 | 0.00202 | -0.0372 | 0.0676 |
| rs548897 | G | A | -0.00966 | 0.00171 | -0.0473 | 0.0563 |
| rs564037 | T | C | 0.01161 | 0.00177 | -0.0267 | 0.0579 |
| rs580016 | C | T | 0.01339 | 0.00229 | 0.0334 | 0.0765 |
| rs6065080 | C | T | 0.01268 | 0.00177 | 0.0425 | 0.0594 |
| rs6093693 | T | C | -0.01032 | 0.00184 | -0.0174 | 0.0612 |
| rs6137567 | C | T | 0.00942 | 0.00172 | -0.0309 | 0.0582 |
| rs613872 | G | T | 0.0175 | 0.00227 | 0.0346 | 0.0743 |
| rs622169 | C | T | -0.00999 | 0.00178 | 0.0719 | 0.0583 |
| rs629681 | T | C | 0.00948 | 0.00172 | 0.0134 | 0.0565 |
| rs639359 | T | G | -0.02374 | 0.00198 | 0.0068 | 0.0665 |
| rs6435670 | T | G | -0.01464 | 0.0019 | 0.0604 | 0.0625 |
| rs6480234 | T | C | -0.00999 | 0.0017 | -0.0649 | 0.0576 |
| rs6485153 | T | C | -0.01129 | 0.00201 | -0.1774 | 0.0668 |
| rs6493265 | C | T | 0.01385 | 0.00174 | 0.0417 | 0.0581 |
| rs6506735 | A | G | 0.01296 | 0.00186 | 0.0959 | 0.063 |
| rs6539284 | T | C | -0.01083 | 0.00173 | 0.0018 | 0.0576 |
| rs6545863 | C | T | -0.01573 | 0.00242 | -0.1688 | 0.0818 |
| rs6556982 | T | G | -0.01151 | 0.00173 | -0.1151 | 0.0842 |
| rs6557171 | C | T | 0.01567 | 0.00181 | -0.1568 | 0.0599 |
| rs6573553 | G | T | -0.01107 | 0.00174 | -0.0178 | 0.057 |
| rs6580699 | T | G | -0.01265 | 0.00173 | -0.1121 | 0.0567 |
| rs6589386 | C | T | -0.00949 | 0.00172 | -0.0099 | 0.0585 |
| rs6672527 | T | C | -0.0101 | 0.00181 | -0.1769 | 0.0618 |
| rs6713957 | A | G | 0.0101 | 0.00178 | 0.111 | 0.0593 |
| rs672644 | T | C | -0.01136 | 0.0019 | -0.0812 | 0.0638 |
| rs6747792 | G | T | 0.01332 | 0.00215 | 0.04 | 0.0708 |
| rs6752812 | A | C | 0.01086 | 0.00184 | -0.089 | 0.0621 |
| rs6805241 | C | T | -0.01413 | 0.00203 | -0.0796 | 0.0677 |
| rs6834271 | G | T | -0.01369 | 0.00235 | 0.081 | 0.0845 |
| rs6949513 | A | G | 0.01104 | 0.00171 | -0.0593 | 0.0576 |
| rs698724 | G | T | 0.01228 | 0.00219 | -0.0464 | 0.0736 |
| rs7085239 | G | A | 0.01451 | 0.00197 | -0.0534 | 0.0668 |
| rs710630 | C | T | -0.01043 | 0.00178 | -0.2658 | 0.0852 |
| rs7119426 | A | G | 0.01173 | 0.00191 | -0.039 | 0.0642 |
| rs7123876 | T | C | 0.01572 | 0.00195 | 0.0649 | 0.0667 |
| rs7185291 | T | C | -0.01049 | 0.00176 | 0.0417 | 0.058 |
| rs7189819 | T | C | 0.01046 | 0.00185 | -0.116 | 0.0607 |
| rs7195278 | T | G | 0.01146 | 0.00173 | -0.0029 | 0.0578 |
| rs7233920 | A | G | -0.01315 | 0.00202 | 0.0976 | 0.066 |
| rs7257460 | T | C | 0.01145 | 0.00189 | -0.149 | 0.062 |
| rs7271519 | T | C | 0.01164 | 0.00185 | 0.0176 | 0.062 |
| rs728054 | G | A | 0.01274 | 0.00177 | 0.0211 | 0.0606 |
| rs730384 | G | A | -0.01016 | 0.00171 | 0.0064 | 0.0567 |
| rs7304782 | A | G | -0.02165 | 0.00197 | 0.1106 | 0.0938 |
| rs7309 | G | A | 0.01658 | 0.0017 | 0.0535 | 0.0572 |
| rs7481514 | G | A | 0.01072 | 0.00178 | -0.0544 | 0.0594 |
| rs7534836 | C | T | -0.00952 | 0.0017 | -0.0015 | 0.0565 |
| rs7543462 | A | G | -0.00973 | 0.00171 | -0.0185 | 0.0559 |
| rs7546833 | A | G | 0.01278 | 0.00217 | 0.0947 | 0.0725 |
| rs7560871 | A | G | -0.01847 | 0.00324 | 0.0633 | 0.1073 |
| rs756912 | C | T | 0.01443 | 0.0017 | -0.0358 | 0.0577 |
| rs7597126 | C | T | 0.01009 | 0.00172 | 0.032 | 0.0569 |
| rs7601713 | A | C | 0.01121 | 0.00194 | 0.0306 | 0.0639 |
| rs7625428 | C | T | -0.00989 | 0.00174 | 0.0204 | 0.0573 |
| rs7685520 | T | C | -0.01045 | 0.00176 | 0.0181 | 0.0593 |
| rs7702649 | C | A | -0.01364 | 0.00219 | 0.093 | 0.0766 |
| rs7723097 | A | C | -0.01023 | 0.00172 | -0.042 | 0.0573 |
| rs7768089 | A | G | 0.00977 | 0.00178 | -0.024 | 0.0588 |
| rs7769945 | G | A | 0.01213 | 0.00219 | -0.069 | 0.0722 |
| rs7803932 | A | G | 0.0143 | 0.00226 | -0.0283 | 0.0755 |
| rs7814699 | G | A | -0.01289 | 0.00234 | -0.0008 | 0.0808 |
| rs7857610 | C | T | -0.0129 | 0.0023 | -0.159 | 0.0754 |
| rs7870597 | T | C | -0.01248 | 0.00188 | 0.0577 | 0.064 |
| rs790647 | C | A | 0.01482 | 0.00202 | -0.1441 | 0.0671 |
| rs7906788 | A | G | 0.00987 | 0.00172 | -0.0522 | 0.0568 |
| rs7910462 | G | A | -0.01215 | 0.00214 | 0.0707 | 0.0746 |
| rs7947194 | A | G | -0.01035 | 0.00176 | 0.0032 | 0.0587 |
| rs7977614 | G | A | 0.01325 | 0.00198 | -0.1475 | 0.0655 |
| rs7981424 | G | A | -0.01147 | 0.00188 | 0.055 | 0.0652 |
| rs7998344 | T | C | -0.01146 | 0.00189 | -0.002 | 0.0626 |
| rs8013500 | T | C | -0.01182 | 0.00213 | 0.0264 | 0.1029 |
| rs8014053 | G | T | 0.0156 | 0.002 | -0.1023 | 0.0676 |
| rs802443 | T | G | -0.00995 | 0.00176 | -0.0574 | 0.0587 |
| rs8058057 | T | C | -0.01339 | 0.00205 | 0.052 | 0.0974 |
| rs8086807 | G | A | 0.01048 | 0.00172 | 0.0718 | 0.057 |
| rs820953 | A | G | 0.01005 | 0.0018 | 0.0428 | 0.0606 |
| rs834143 | T | C | -0.01015 | 0.0017 | 0.0596 | 0.0564 |
| rs891389 | C | T | 0.01466 | 0.00178 | 0.0412 | 0.0591 |
| rs893363 | A | G | 0.01008 | 0.00174 | -0.141 | 0.059 |
| rs895606 | G | A | -0.01204 | 0.0017 | 0.0048 | 0.0576 |
| rs912069 | C | T | -0.01209 | 0.00208 | -0.1173 | 0.0698 |
| rs9285211 | G | T | -0.01643 | 0.00299 | -0.0914 | 0.0995 |
| rs9303690 | C | T | 0.01316 | 0.00227 | -0.1131 | 0.0736 |
| rs9311952 | C | A | -0.013 | 0.00237 | 0.0295 | 0.0771 |
| rs9320493 | G | A | -0.01394 | 0.0024 | 0.0133 | 0.1153 |
| rs9324593 | G | A | -0.01633 | 0.00246 | -0.0995 | 0.0832 |
| rs9386110 | T | C | -0.0163 | 0.00272 | -0.0227 | 0.1219 |
| rs9436866 | C | A | 0.01882 | 0.00289 | 0.0221 | 0.0957 |
| rs950493 | T | C | -0.0128 | 0.00233 | 0.027 | 0.0774 |
| rs952530 | T | C | 0.01003 | 0.00172 | 0.0708 | 0.0582 |
| rs9557378 | A | G | -0.01094 | 0.00193 | 0.0834 | 0.0633 |
| rs9647518 | G | A | 0.00935 | 0.0017 | 0.0461 | 0.0571 |
| rs967885 | C | T | -0.01011 | 0.00185 | 0.0057 | 0.0611 |
| rs968827 | T | C | -0.01102 | 0.00187 | 0.0121 | 0.0626 |
| rs981883 | G | A | -0.0113 | 0.00174 | -0.0889 | 0.0594 |
| rs9824301 | C | A | 0.01459 | 0.00177 | -0.0387 | 0.0591 |
| rs983798 | G | A | 0.01284 | 0.00208 | -0.095 | 0.0676 |
| rs9846711 | C | T | 0.0113 | 0.00176 | -0.0205 | 0.0586 |
| rs9964724 | C | T | -0.01978 | 0.00183 | -0.0038 | 0.0618 |
| rs998887 | A | C | 0.01272 | 0.0017 | 0.0639 | 0.0567 |

Supplementary Table 4: GWAS summary statistics used to conduct an MR of the effect of intelligence on liability to nsCL/P.

| SNP | Effect Allele | Other Allele | Intelligence Beta | Intelligence SE | nsCL/P Beta | nsCL/P SE |
| --- | --- | --- | --- | --- | --- | --- |
| rs10010325 | C | A | -0.01956 | 0.00285 | 0.0591 | 0.0573 |
| rs1007934 | G | A | -0.018 | 0.00289 | -0.0168 | 0.0576 |
| rs10089969 | T | G | -0.02318 | 0.00341 | -0.0649 | 0.0681 |
| rs10129426 | G | A | -0.01926 | 0.00287 | -0.0066 | 0.058 |
| rs10148349 | T | C | -0.01945 | 0.00339 | -0.1366 | 0.0677 |
| rs10180461 | C | T | 0.01949 | 0.00292 | 0.0248 | 0.0589 |
| rs1035738 | T | C | 0.05092 | 0.00769 | -0.0259 | 0.2472 |
| rs1043595 | G | A | -0.02097 | 0.00317 | -0.0154 | 0.0639 |
| rs1044258 | T | C | 0.02533 | 0.00298 | 0.0536 | 0.0586 |
| rs10499812 | G | A | -0.0215 | 0.00325 | -0.0594 | 0.0666 |
| rs10733789 | T | C | -0.01692 | 0.00307 | 0.0019 | 0.0616 |
| rs10875914 | G | A | 0.02373 | 0.00291 | 0.0743 | 0.0572 |
| rs10876864 | G | A | 0.01679 | 0.00289 | 0.1603 | 0.0576 |
| rs10996430 | A | G | -0.02036 | 0.00332 | -0.0765 | 0.0656 |
| rs11012 | C | T | 0.02562 | 0.00369 | -0.1791 | 0.0744 |
| rs11164719 | C | T | 0.01658 | 0.00285 | -0.1026 | 0.0566 |
| rs11186624 | C | A | -0.01791 | 0.00297 | 0.0496 | 0.0605 |
| rs11264680 | C | T | -0.01863 | 0.0031 | 0.1134 | 0.0606 |
| rs1133415 | G | A | -0.01629 | 0.00286 | -0.0031 | 0.0558 |
| rs1144593 | G | A | 0.02276 | 0.00309 | -0.0496 | 0.062 |
| rs1145123 | C | T | -0.01896 | 0.00286 | -0.1208 | 0.0565 |
| rs11649476 | T | C | 0.01712 | 0.00287 | -0.0772 | 0.0562 |
| rs11679484 | A | C | -0.01634 | 0.00296 | -0.0276 | 0.0582 |
| rs11720523 | C | A | -0.01622 | 0.00288 | -0.0411 | 0.0577 |
| rs12032756 | G | A | 0.02728 | 0.00343 | 0.1344 | 0.0701 |
| rs12124523 | C | T | -0.03381 | 0.00458 | -0.1342 | 0.094 |
| rs12325113 | C | T | -0.02564 | 0.00291 | 0.003 | 0.0808 |
| rs12446238 | G | A | -0.01815 | 0.00288 | -0.0461 | 0.0571 |
| rs1245214 | G | A | 0.01893 | 0.0033 | -0.1005 | 0.0669 |
| rs1248853 | G | A | 0.04183 | 0.00734 | -0.0024 | 0.2245 |
| rs12646225 | C | T | -0.02975 | 0.00464 | -0.0583 | 0.0901 |
| rs13010010 | C | T | -0.02401 | 0.00294 | -0.1483 | 0.0591 |
| rs13012916 | G | T | 0.01632 | 0.00294 | -0.0823 | 0.0592 |
| rs13064915 | T | C | 0.01708 | 0.00304 | 0.0469 | 0.0605 |
| rs13107325 | C | T | 0.05434 | 0.00542 | 0.1045 | 0.121 |
| rs1317149 | T | C | -0.0187 | 0.00299 | -0.0449 | 0.083 |
| rs13197574 | C | T | 0.03972 | 0.00453 | 0.2978 | 0.097 |
| rs13253386 | T | G | -0.01836 | 0.00285 | 0.086 | 0.0561 |
| rs1360382 | A | G | -0.02234 | 0.00304 | -0.05 | 0.0609 |
| rs1394466 | A | G | 0.02239 | 0.00285 | -0.0055 | 0.0558 |
| rs1437721 | A | G | 0.01697 | 0.00296 | -0.0975 | 0.0595 |
| rs1473634 | A | G | -0.01906 | 0.00311 | 0.0629 | 0.0621 |
| rs1479679 | C | T | -0.02436 | 0.00426 | -0.1024 | 0.0886 |
| rs1484237 | C | T | 0.01893 | 0.00342 | -0.0858 | 0.0687 |
| rs1507010 | G | A | 0.01728 | 0.00286 | 0.0491 | 0.0564 |
| rs1523048 | T | C | 0.01776 | 0.00296 | 0.1143 | 0.0579 |
| rs1560359 | A | C | -0.01776 | 0.00321 | -0.0223 | 0.0659 |
| rs16874969 | G | A | 0.02082 | 0.00376 | -0.0461 | 0.0752 |
| rs17151739 | A | C | 0.02214 | 0.0031 | -0.051 | 0.0627 |
| rs17692512 | T | G | 0.01945 | 0.00348 | 0.098 | 0.0704 |
| rs17765227 | G | A | 0.0182 | 0.00329 | 0.0181 | 0.0665 |
| rs17825119 | T | G | -0.01709 | 0.00285 | 0.1152 | 0.0788 |
| rs1862499 | G | A | 0.01647 | 0.003 | 0.0577 | 0.0592 |
| rs1906252 | C | A | -0.03071 | 0.00285 | 0.116 | 0.0568 |
| rs1908268 | A | G | 0.01818 | 0.00285 | -0.0945 | 0.0565 |
| rs1979611 | T | C | -0.0157 | 0.00285 | 0.0828 | 0.0575 |
| rs2012512 | A | G | -0.01569 | 0.00286 | 0.0761 | 0.0573 |
| rs2052572 | A | G | -0.0184 | 0.00304 | -0.0545 | 0.06 |
| rs2054122 | T | G | 0.01681 | 0.00286 | 0.0255 | 0.0577 |
| rs2074103 | C | A | -0.01843 | 0.00287 | -0.0941 | 0.0566 |
| rs2143103 | A | G | 0.02382 | 0.00415 | -0.0917 | 0.0875 |
| rs2235117 | C | T | 0.02069 | 0.00362 | -0.0001 | 0.0697 |
| rs2289328 | A | G | -0.02241 | 0.00385 | -0.0022 | 0.0766 |
| rs2294405 | G | A | -0.01712 | 0.00303 | 0.0411 | 0.0607 |
| rs2302210 | G | A | 0.02508 | 0.00398 | -0.0477 | 0.1191 |
| rs2333611 | A | G | 0.01606 | 0.00294 | 0.0955 | 0.0592 |
| rs2343606 | T | C | 0.02021 | 0.00321 | -0.1344 | 0.0641 |
| rs243071 | A | G | -0.01812 | 0.00289 | 0.1387 | 0.0571 |
| rs2436391 | C | T | -0.02178 | 0.00286 | 0.0556 | 0.0575 |
| rs2478288 | T | C | -0.02263 | 0.00326 | 0.0466 | 0.065 |
| rs2545796 | C | T | -0.01793 | 0.00287 | -0.0898 | 0.0793 |
| rs276626 | G | A | -0.02075 | 0.0038 | -0.0964 | 0.0761 |
| rs2862954 | C | T | 0.01669 | 0.00286 | -0.0078 | 0.0573 |
| rs2989868 | A | G | 0.01822 | 0.00285 | -0.1066 | 0.0563 |
| rs3741434 | C | T | -0.02242 | 0.0041 | 0.1238 | 0.08 |
| rs3748256 | G | A | 0.01929 | 0.00293 | 0.021 | 0.0585 |
| rs3749034 | G | A | 0.01995 | 0.00345 | 0.0802 | 0.0699 |
| rs3755496 | G | T | -0.02916 | 0.00517 | 0.0079 | 0.1026 |
| rs3757966 | A | G | -0.0158 | 0.00286 | 0.0056 | 0.0574 |
| rs3850610 | C | A | -0.02114 | 0.00332 | 0.0793 | 0.0651 |
| rs3943667 | T | C | -0.0185 | 0.00313 | 0.0272 | 0.0883 |
| rs4129585 | A | C | -0.01931 | 0.00288 | 0.0027 | 0.0577 |
| rs4244210 | G | A | -0.01658 | 0.00302 | -0.0026 | 0.0599 |
| rs4261436 | C | T | -0.02017 | 0.00288 | 0.0297 | 0.0569 |
| rs4266584 | A | C | -0.0171 | 0.00306 | -0.094 | 0.0615 |
| rs4342312 | C | A | 0.01793 | 0.00294 | -0.0191 | 0.0584 |
| rs4438512 | A | G | 0.01714 | 0.00288 | -0.0041 | 0.0575 |
| rs4664442 | A | G | -0.02041 | 0.00286 | -0.0203 | 0.0568 |
| rs4696283 | G | A | -0.01936 | 0.00285 | 0.0445 | 0.0568 |
| rs4744250 | A | G | 0.01834 | 0.003 | 0.044 | 0.0827 |
| rs4788585 | G | A | 0.0188 | 0.00323 | 0.0337 | 0.0628 |
| rs4949455 | G | A | -0.02123 | 0.00295 | -0.0148 | 0.0592 |
| rs501299 | A | C | -0.01642 | 0.00294 | -0.0703 | 0.0591 |
| rs538827 | T | C | 0.01905 | 0.00304 | -0.0775 | 0.0603 |
| rs6047982 | T | G | -0.01715 | 0.00293 | 0.1198 | 0.0816 |
| rs6066968 | A | G | 0.02588 | 0.0029 | -0.0259 | 0.0581 |
| rs633158 | C | T | -0.01863 | 0.00312 | 0.0121 | 0.0626 |
| rs6539284 | T | C | -0.02243 | 0.00291 | 0.0018 | 0.0576 |
| rs658912 | G | A | 0.01667 | 0.00294 | 0.0452 | 0.059 |
| rs6674176 | A | G | 0.0236 | 0.00377 | 0.1731 | 0.0741 |
| rs6678734 | A | G | 0.02342 | 0.00285 | 0.009 | 0.0579 |
| rs6744254 | T | C | 0.0158 | 0.00285 | 0.012 | 0.0562 |
| rs6802890 | A | G | 0.03099 | 0.00285 | -0.0371 | 0.0566 |
| rs6882046 | G | A | 0.02586 | 0.00321 | -0.0963 | 0.0637 |
| rs6903716 | G | A | -0.01842 | 0.00314 | 0.0536 | 0.062 |
| rs6911407 | C | A | 0.028 | 0.00295 | -0.0439 | 0.0581 |
| rs6930924 | A | G | 0.02566 | 0.00467 | 0.0197 | 0.0959 |
| rs6975134 | C | T | 0.02194 | 0.00291 | 0.0683 | 0.0586 |
| rs6994361 | C | T | 0.01821 | 0.00292 | 0.0676 | 0.0582 |
| rs7128489 | C | T | 0.01967 | 0.00326 | -0.0336 | 0.0649 |
| rs7189819 | T | C | 0.02509 | 0.00306 | -0.116 | 0.0607 |
| rs7245 | A | G | 0.02093 | 0.00286 | 0.0511 | 0.0798 |
| rs730934 | C | T | -0.01782 | 0.00286 | 0.0563 | 0.0574 |
| rs7431278 | C | T | -0.02469 | 0.00305 | -0.1024 | 0.0839 |
| rs7502208 | G | A | -0.01772 | 0.00295 | 0.1079 | 0.0581 |
| rs756912 | C | T | 0.01954 | 0.00285 | -0.0358 | 0.0577 |
| rs7577045 | G | A | -0.0174 | 0.00288 | 0.03 | 0.0569 |
| rs7626560 | C | T | -0.02393 | 0.00389 | -0.0729 | 0.0782 |
| rs768851 | G | A | 0.01753 | 0.00294 | 0.0512 | 0.0581 |
| rs7871404 | A | G | -0.02301 | 0.00364 | -0.0866 | 0.0723 |
| rs7942897 | C | T | -0.04113 | 0.00689 | -0.2221 | 0.1397 |
| rs799447 | T | G | 0.02082 | 0.00305 | -0.0534 | 0.0621 |
| rs830383 | A | G | 0.01952 | 0.00295 | -0.0404 | 0.0584 |
| rs875361 | A | G | -0.01623 | 0.00287 | -0.099 | 0.0572 |
| rs909674 | C | A | -0.02219 | 0.00328 | -0.0328 | 0.0646 |
| rs913264 | C | T | -0.01881 | 0.00312 | -0.07 | 0.0618 |
| rs9287826 | C | A | -0.01906 | 0.00333 | 0.0418 | 0.0654 |
| rs929456 | T | G | 0.01618 | 0.00295 | 0.0744 | 0.0597 |
| rs9424981 | T | C | 0.03332 | 0.00353 | -0.062 | 0.0939 |
| rs9436866 | C | A | 0.03309 | 0.00481 | 0.0221 | 0.0957 |
| rs946468 | G | A | -0.02227 | 0.00359 | 0.0139 | 0.0981 |
| rs958217 | A | G | 0.01776 | 0.00304 | -0.0988 | 0.06 |

Supplementary Table 5: Causal effect estimates of genetic liability for nsCL/P on educational attainment derived by the primary MR analysis (IVW) as well as sensitivity analyses.

| Method | N SNPs | B | se | pval | 95% C.I. |
| --- | --- | --- | --- | --- | --- |
| IVW | 12 | 0.002 | 0.003 | 0.417 | -0.003- 0.007 |
| MR Egger | 12 | -0.008 | 0.006 | 0.226 | -0.019- 0.004 |
| Weighted median | 12 | 0.001 | 0.002 | 0.496 | -0.003- 0.005 |
| Weighted mode | 12 | -0.0003 | 0.002 | 0.906 | -0.003- 0.005 |

Supplementary Table 6: Causal effect estimates of genetic liability for nsCL/P on intelligence derived by the primary MR analysis (IVW) as well as sensitivity analyses.

| Method | N SNPs | B | se | pval | 95% C.I. |
| --- | --- | --- | --- | --- | --- |
| IVW | 12 | 0.002 | 0.006 | 0.669 | -0.009- 0.014 |
| MR Egger | 12 | -0.018 | 0.014 | 0.229 | -0.045- 0.009 |
| Weighted median | 12 | -0.002 | 0.004 | 0.507 | -0.010- 0.005 |
| Weighted mode | 12 | -0.004 | 0.004 | 0.333 | -0.011- 0.003 |

Supplementary Table 7: Causal effect estimates of genetic liability for nsCL/P on educational attainment derived by the primary MR analysis (IVW) as well as sensitivity analyses. [RESTRICTING TO SNPs FROM EUROPEAN nsCL/P GWAS].

| Method | N SNPs | B | se | pval | 95% C.I. |
| --- | --- | --- | --- | --- | --- |
| IVW | 6 | -0.0002 | 0.004 | 0.949 | -0.008- 0.007 |
| MR Egger | 6 | -0.0061 | 0.011 | 0.596 | -0.027- 0.015 |
| Weighted median | 6 | 0.0001 | 0.002 | 0.953 | -0.005- 0.004 |
| Weighted mode | 6 | -0.0003 | 0.002 | 0.886 | -0.004- 0.004 |

Supplementary Table 8: Causal effect estimates of genetic liability for nsCL/P on intelligence derived by the primary MR analysis (IVW) as well as sensitivity analyses. [RESTRICTING TO SNPs FROM EUROPEAN nsCL/P GWAS].

| Method | N SNPs | B | se | pval | 95% C.I. |
| --- | --- | --- | --- | --- | --- |
| IVW | 6 | -0.003 | 0.004 | 0.531 | -0.010- 0.005 |
| MR Egger | 6 | -0.013 | 0.010 | 0.275 | -0.034- 0.007 |
| Weighted median | 6 | -0.003 | 0.004 | 0.453 | -0.010- 0.005 |
| Weighted mode | 6 | -0.007 | 0.004 | 0.143 | -0.015- 0.0009 |

Supplementary Table 9: Causal effect estimates of educational attainment on nsCL/P derived by the primary MR analysis (IVW) as well as sensitivity analyses.

| Method | N SNPs | OR | pval | 95% CI |
| --- | --- | --- | --- | --- |
| IVW | 368 | 1.337 | 0.323 | 0.752- 2.380 |
| MR Egger | 368 | 0.464 | 0.562 | 0.035- 6.229 |
| Weighted Median | 368 | 1.153 | 0.726 | 0.496- 2.678 |
| Weighted Mode | 368 | 1.142 | 0.926 | 0.061- 21.393 |

Supplementary Table 10: Causal effect estimates of intelligence on nsCL/P derived by the primary MR analysis (IVW) as well as sensitivity analyses.

| Method | N SNPs | OR | pval | 95% CI |
| --- | --- | --- | --- | --- |
| IVW | 132 | 1.062 | 0.851 | 0.567- 1.987 |
| MR Egger | 132 | 5.662 | 0.264 | 0.274- 117.036 |
| Weighted Median | 132 | 0.932 | 0.865 | 0.410- 2.114 |
| Weighted Mode | 132 | 0.783 | 0.842 | 0.007- 8.751 |

**CODE**

C1. LDScore Regression analysis- Estimation of the genetic correlation among nsCL/P and educational attainment.

*#Formatting the Cleft GWAS file to be readable by the LDSC package#*

```
./munge_sumstats.py \
 --sumstats BonnTDT.MA.PRSice.txt \
 --N 3987 \
 --out nsCL \
 --merge-alleles w_hm3.snplist
```

*#Formatting the EA GWAS file to be readable by the LDSC package#*

```
./munge_sumstats.py \
 --sumstats GWAS_EA_excl23andMe.txt \
 --N 766345 \
 --out EA_GWAS \
 --merge-alleles w_hm3.snplist
```

#Genetic Correlation Analysis#

```
./ldsc.py \
 --rg nsCL.sumstats.gz,EA_GWAS.sumstats.gz \
 --ref-ld-chr eur_w_ld_chr/ \
 --w-ld-chr eur_w_ld_chr/ \
 --out nsCL_EA
```

C2. LDScore Regression analysis- Estimation of the genetic correlation among nsCL/P and intelligence.

*#Formatting the Cleft GWAS file to be readable by the LDSC package#*

```
./munge_sumstats.py \
 --sumstats BonnTDT.MA.PRSice.txt \
 --N 3987 \
 --out nsCL \
 --merge-alleles w_hm3.snplist
```

*#Formatting the CP GWAS file to be readable by the LDSC package#*

```
./munge_sumstats.py \
 --sumstats GWAS_CP_all.txt \
 --N 257828 \
 --out CP_GWAS \
 --merge-alleles w_hm3.snplist
```

#Genetic Correlation Analysis#

./ldsc.py \

--rg nsCL.sumstats.gz,CP_GWAS.sumstats.gz \

--ref-ld-chr eur_w_ld_chr/ \

--w-ld-chr eur_w_ld_chr/ \

--out nsCL_CP

C3. Two Sample MR analysis- The causal effect of nsCL/P liability on Educational attainment.

*#Obtaining instruments from the Cleft GWAS#*

cleft_instr<- subset(BonnTDT.MA.PRSice, SNP %in% c("rs560426", "rs861020", "rs987525", "rs7078160", "rs227731", "rs13041247", "rs742071", "rs7590268", "rs7632427", "rs12543318", "rs8001641", "rs1873147"))

*#Formatting the file to be readable by the MRBase package#*

cleft_instr<- format_data(cleft_instr, type = "exposure", snps = NULL, header = TRUE, phenotype_col = "nsCLP", snp_col = "SNP", beta_col = "BETA", se_col = "SE", effect_allele_col = "A1", other_allele_col = "A2", pval_col = "P")

*#Clumping the instruments r^2= 0.01#*

cleft_instr<- clump_data(cleft_instr, clump_r2 = 0.01)

*#Extracting the instruments' effect sizes from the Educational Attainment GWAS#*

EA_out<- read_outcome_data(snps = cleft_instr$SNP, filename = "Lee_ed_data,csv", sep = ",", snp_col = "SNP", beta_col = "beta.outcome", se_col = "se.outcome", effect_allele_col = "effect_allele.outcome", other_allele_col = "other_allele.outcome", pval_col = "pval.outcome")

*#Harmonising exposure- outcome#*

cleft_EA<- harmonise_data(exposure_dat = cleft_instr, outcome_dat = EA_out)

*#Analysis#*

res_Cleft_EA<- mr(cleft_EA, method_list=c("mr_egger_regression", "mr_ivw", "mr_weighted_mode", "mr_weighted_median"))

C4. Two Sample MR analysis- The causal effect of nsCL/P liability on Intelligence.

*#Obtaining instruments from the Cleft GWAS#*

cleft_instr<- subset(BonnTDT.MA.PRSice, SNP %in% c("rs560426", "rs861020", "rs987525", "rs7078160", "rs227731", "rs13041247", "rs742071", "rs7590268", "rs7632427", "rs12543318", "rs8001641", "rs1873147"))

*#Formatting the file to be readable by the MRBase package#*

cleft_instr<- format_data(cleft_instr, type = "exposure", snps = NULL, header = TRUE, phenotype_col = "nsCLP", snp_col = "SNP", beta_col = "BETA", se_col = "SE", effect_allele_col = "A1", other_allele_col = "A2", pval_col = "P")

*#Clumping the instruments r^2= 0.01#*

cleft_instr<- clump_data(cleft_instr, clump_r2 = 0.01)

*#Extracting the instruments' effect sizes from the Intelligence GWAS#*

CP_out<- read_outcome_data(snps = cleft_instr$SNP, filename = "Lee_iq_data,csv", sep = ",", snp_col = "SNP", beta_col = "beta.outcome", se_col = "se.outcome", effect_allele_col = "effect_allele.outcome", other_allele_col = "other_allele.outcome", pval_col = "pval.outcome")

*#Harmonising exposure- outcome#*

cleft_CP<- harmonise_data(exposure_dat = cleft_instr, outcome_dat = CP_out)

*#Analysis#*

res_Cleft_CP<- mr(cleft_CP, method_list=c("mr_egger_regression", "mr_ivw", "mr_weighted_mode", "mr_weighted_median"))

C5. Two Sample MR analysis- The causal effect of Educational Attainment on nsCL/P.

*#Restricting the EA GWAS SNPs to SNPs present in the Cleft GWAS#*

EA_gwas_results_common<-GWAS_EA_excl23andMe[GWAS_EA_excl23andMe$MarkerName %in% BonnTDT.MA.PRSice$SNP,]

*#Formatting the data to be readable by MRBase Package#*

EA_common<- format_data(EA_gwas_results_common, type = "exposure", snps = NULL, header = TRUE, snp_col = "MarkerName", beta_col = "Beta", se_col = "SE", eaf_col = "EAF", effect_allele_col = "A1", other_allele_col = "A2", pval_col = "Pval")

*#Defining the instruments#*

EA_instr<- EA_common [which (EA_common$pval.exposure < 5e-08), ]

EA_instr<- EA_instr [which (EA_instr$eaf.exposure >= 0.01), ]

EA_instr<- clump_data(EA_instr, clump_r2 = 0.01)

*#Extracting the instruments' effect sizes from the Cleft GWAS#*

cleft_out<- read_outcome_data(snps = EA_instr$SNP, filename = "Cleft_outcome,csv", sep = ",", snp_col = "SNP", beta_col = "beta.outcome", se_col = "se.outcome", effect_allele_col = "effect_allele.outcome", other_allele_col = "other_allele.outcome", pval_col = "pval.outcome")

*#Harmonising exposure- outcome#*

EA_clf<- harmonise_data(EA_instr, cleft_out)

*#Analysis#*

res_EA_cleft <- mr(EA_clf, method_list=c("mr_egger_regression", "mr_ivw", "mr_weighted_median", "mr_weighted_mode"))

C6. Two Sample MR analysis- The causal effect of Intelligence on nsCL/P.

*#Restricting the EA GWAS SNPs to SNPs present in the Cleft GWAS#*

CP_gwas_results_common<-GWAS_CP_all[GWAS_CP_all$MarkerName %in% BonnTDT.MA.PRSice$SNP,]

*#Formatting the data to be readable by MRBase Package#*

CP_common<- format_data(CP_gwas_results_common, type = "exposure", snps = NULL, header = TRUE, snp_col = "MarkerName", beta_col = "Beta", se_col = "SE", eaf_col = "EAF", effect_allele_col = "A1", other_allele_col = "A2", pval_col = "Pval")

*#Defining the instruments#*

CP_instr<- CP_common [which (CP_common$pval.exposure < 5e-08), ]

CP_instr<- CP_instr [which (CP_instr$eaf.exposure >= 0.01), ]

CP_instr<- clump_data(CP_instr, clump_r2 = 0.01)

*#Extracting the instruments' effect sizes from the Cleft GWAS#*

cleft_out<- read_outcome_data(snps = CP_instr$SNP, filename = "Cleft_outcome,csv", sep = ",", snp_col = "SNP", beta_col = "beta.outcome", se_col = "se.outcome", effect_allele_col = "effect_allele.outcome", other_allele_col = "other_allele.outcome", pval_col = "pval.outcome")

*#Harmonising exposure- outcome#*

CP_clf<- harmonise_data(CP_instr, cleft_out)

*#Analysis#*

res_CP_cleft <- mr(CP_clf, method_list=c("mr_egger_regression", "mr_ivw", "mr_weighted_median", "mr_weighted_mode"))
